# X-chromosome inactivation and its implications for human disease

**DOI:** 10.1101/076950

**Authors:** Joost Gribnau, Tahsin Stefan Barakat

**Affiliations:** Department of Developmental Biology, Erasmus MC – University Medical Center, Rotterdam, The Netherlands; MRC Center for Regenerative Medicine, Institute for Stem Cell Research, School of Biological Sciences, University of Edinburgh, Edinburgh, United Kingdom

**Keywords:** Clinical Genetics, X-chromosome inactivation disease implications, skewing, manifesting heterozygotes, mosaicism

## Abstract

In humans and other mammals, female cells carry two X-chromosomes, whereas male cells carry a single X and Y-chromosome. To achieve an equal expression level of X-linked genes in both sexes, a dosage compensation mechanism evolved, which results in transcriptional silencing of one X-chromosome in females. X chromosome inactivation (XCI) is random with respect to the parental origin of the X, occurs early during embryonic development, and is then stably maintained through a near infinite number of cell divisions. As a result of this, every female individual consists of a mosaic of two different cell populations, in which either the maternally or paternally derived X-chromosome is inactivated. As the X-chromosome harbors more than a thousand genes, of which many are implicated in human disease when mutated, this mosaicism has important disease implications. Whereas X-linked disorders are usually more severe in hemizygous males harboring a single X-chromosome, a more variable phenotype is observed in females. This variability is a direct consequence of the XCI-mosaicism, and is affected by the randomness of the XCI process. Here we review the latest insights into the regulation of this important female specific process, and discuss mechanisms that influence mosaicism in females, with a focus on the clinical consequences related to X-linked diseases in females.

## X-CHROMOSOME INACTIVATION AND THE NEED FOR DOSAGE COMPENSATION

Both sexes contain an equal number of autosomal chromosomes, and a balanced dosage of X-linked genes is needed to accomplish functional cell physiology. In heterogametic species, the evolution of a single gene-rich X and gene-pore Y-chromosome in males results in potential dosage problems, as in males only a single X-chromosome is responsive for the same functions as two X-chromosomes in females. Many genes act in dose-dependent manners, and therefore during the attrition of the proto-Y, monosomy of the proto-X-chromosome in males is believed to be compensated by an up-regulation of gene transcription from the developing X-chromosome, thereby restoring the equilibrium between autosomal and X-linked genes^1^. For females, this up-regulation of X-linked gene transcription would further disrupt the desired equilibrium, as the presence of two, highly transcribed, X-chromosomes would disrupt equal gene dosage between the autosomes and sex-chromosomes. To prevent this potential problem, in placental mammals such as humans, dosage compensation of X-linked genes between both sexes is achieved by inactivation of one of the two X-chromosomes in females, a process called X-chromosome inactivation (XCI)^1,2^. XCI occurs during early female development, and results in heterochromatinization and silencing of the X-chromosome, which is maintained during subsequent cell divisions throughout life^3^. XCI leads to mono-allelic expression of most X-linked genes, with the exception of genes located in the pseudo-autosomal regions (PARs) and a number of other genes escaping from inactivation^4^. These latter genes often have X-degenerate counterparts on the Y-chromosome, which also carries a PAR homologous to that of the X-chromosome. Since expression of X-encoded genes from this chromosome and the single X-chromosome in male cells is two-fold up-regulated compared to autosomes, the proper dosage of X-encoded genes is restored, thereby complying to Ohno’s hypothesis^1,5^.

As a consequence of the stable XCI maintenance, adult females are a mosaic of two different cell populations, in which either the maternal or paternal X-chromosome is inactivated. As many X-linked genes can contribute to human disease, this mosaicism is an important modifier of X-linked genetic disorders. Whereas male patients suffering from X-linked mutations are usually strongly affected, the phenotype of X-linked disorders in females is highly variable. The origin of this variability is caused by the XCI-process. Here, we will first discuss the latest insights into the regulation of the XCI-process, followed by a detailed discussion on the mechanisms contributing to variability of X-linked disorders in females.

### Lessons from mouse ES cells for the regulation of XCI

In mice, XCI occurs in two different forms^6^’^7^. Around the 2-to 8-cell stage, imprinted-XCI is initiated, in which the paternal X-chromosome is silenced. Imprinted-XCI is maintained in extra-embryonic tissues, but is reversed in the epiblast which will form the embryo, around day 4.25 (E4.25) of development. Then, a second wave of random-XCI, in which either the maternal or paternal X-chromosome is silenced, is initiated around E5.5 post-implantation, and this process is finalized approximately a day later. As mouse embryonic stem (ES) cells can be derived from the preimplantation epiblast, when both X-chromosomes are active, these cells can be used as an ex vivo model to study the XCI-process, as they undergo XCI upon differentiation^7,8^.

Early studies in mouse and man have shown that a region on the X-chromosome, called the X-inactivation-center (XIC) is crucial for XCI to occur (**Figure 1**). Within the XIC-region, several genes have been localized which execute XCI. The most important gene is the X-inactive-specific-transcript (Xist in mouse, XIST in human), which encodes a non-coding RNA molecule^9–11^. This RNA becomes upregulated upon differentiation, is able to spread along the future inactive X-chromosome (Xi) and attracts chromatin modifying enzymes which establish XCI-silencing. A major focus on XCI-research has been the regulation of Xist-expression, and the question why always one X-chromosome is kept active per diploid genome. In mouse, XCI appears to be directed by a stochastic mechanism, where XCI-initiation is regulated by autosomally-encoded XCI-inhibitors, and X-linked XCI-activators^12^. As the latter are X-linked, they will be expressed in a higher dose in females compared to males, and can explain female-specific XCI-initiation. One such factor identified is Rnf12, which is located in close proximity to Xist^13–15^. RNF12 is an E3 ubiquitin ligase, which is able to target the ES cell factor and XCI-inhibitor REX1 for proteasomal degradation^16^. REX1 represses Xist directly but also stimulates expression of Tsix^16,17^, which is an antisense transcript being transcribed throughout Xist, thereby inhibiting Xist-expression^18^. Hence, when Rnf12 becomes upregulated in female cells upon development, the break on Xist-expression is released, allowing female-specific Xist-expression. The latter is only possible when the Xist locus is located in a transcription prone environment, and as such there is an important role for the cis-acting elements and genes surrounding the Xist-locus, including the cis-acting genes Jpx, Ftx and the Xpr region^19^. Also Tsix is regulated by a wide range of genes and elements located in its vicinity^20^, and together, these elements and genes located on both sides of Xist and Tsix might play an important role in determining which of the two X-chromosomes is more likely to become inactivated. Whereas the role of Rnf12 in the regulation of XCI in ES cells and in imprinted-XCI is well established^13-15^, its role in random-XCI in the mouse epiblast has remained a matter of debate^21^. So far, most studies have focused on mouse XCI, and therefore much less is known about the timing and factors directing human XCI. In human, TSIX is only partially conserved and does not overlap with XIST, and a role for RNF12 in human XCI has yet to be confirmed. Unfortunately, most human ES cells are in a post-XCI state which prevents investigations on XCI-initiation^22^. Few studies have addressed XCI-initiation in human embryos, with conflicting results, where one study found XCI-initiation in cleavage stage embryos comparable to the mouse^23^, whereas another study found XCI-initiation on both X-chromosomes at the blastocyst stage without silencing, which only occurs at later stages on a single X-chromosome^24^. A recent study has used *in vitro* culture of human embryos on decidualized endometrial stromal cells up to E8 of development, and found H3K27me3 on one X-chromosome in cells from trophectoderm and hypoblast indicative for XCI, but not in epiblast cells, arguing that XCI during human development might be regulated in a lineage specific fashion^25^. Also, it has remained a matter of debate whether imprinted-XCI can occur in human^26,27^. Future research is needed to address many of these open questions.

**Figure 1.**
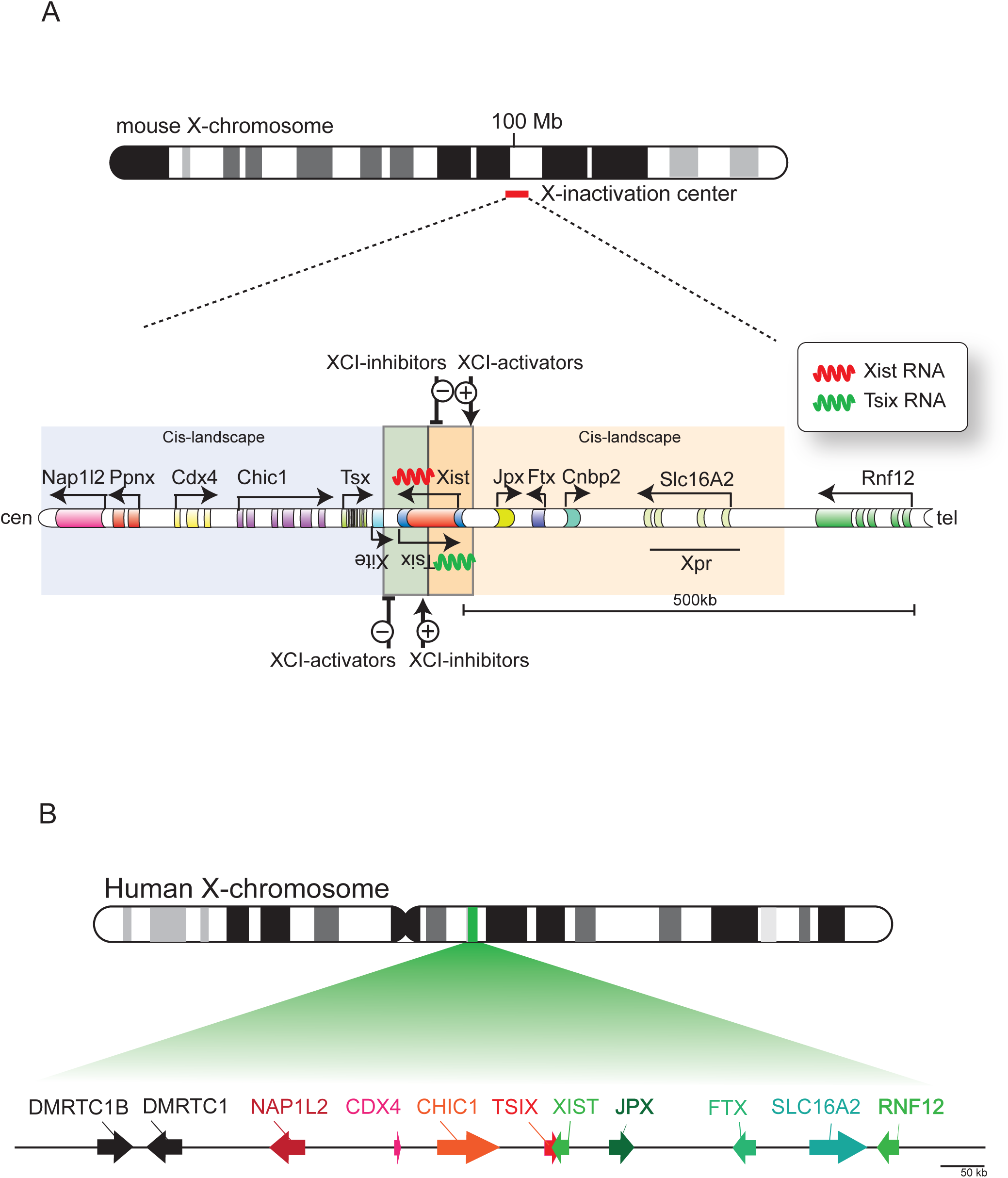
The X-inactivation center and genes regulating X-chromosome inactivation. A) The X-inactivation center in mouse contains the non-coding gene Xist and its antisense partner Tsix. Transcription trough the Tsix locus can inhibit Xist expression, and Xist RNA, when upregulated, can coat the future inactive X-chromosome and recruits chromatin modifying enzymes, which results in gene silencing. Both genes are regulated by *trans-*acting autosomally-encoded XCI-inhibitors and X-linked XCI-activators, and a *cis-*acting landscape, which together ensure female specific initiation of the important X-chromosome inactivation process. B) The homologous X-inactivation center in human

## MENDELIAN INHERITANCE OF X-LINKED TRAITS

Many decades ago, prior to the discovery of chromosomes and genes, it was noticed that several diseases, including color-blindness and hemophilia, were occurring almost only in males (reviewed in^28^). It was noticed that these diseases were not transmitted from affected males to their sons, but that healthy daughters could give birth to sons who were again affected. Although at that time no explanation existed for this way of inheritance, it is now clear that this pattern of inheritance is typical for X-linked traits. The first X-linked trait, the white-eye characteristic, was discovered more than a century ago in *Drosophila^29^.* It was found that when a white-eye male fly was bred to a red-eye female, all offspring had red eyes. When this offspring was inbred with each other, a quarter of all offspring had again white eyes, and these white-eyed flies were all males. From this it was concluded that the trait for eye color in flies must be sex-linked, and that the red eyes trait in flies is dominant, whereas the white-eye characteristic is recessive^29^. This and other work in flies was extended to the X and Y-system in human, leading to the first rules on X-linked inheritance. These rules can still be found in nowadays textbooks of genetics, and state that a character is dominant when it is expressed in a heterozygote, and recessive when not. X-linked recessive diseases almost exclusively affect males (hemizygous for the X-chromosome), and rarely affect homozygous recessive females. Affected males cannot transmit their trait to their sons, rendering male-to-male transmission impossible. Transmission however can occur through their daughters, since all daughters will be obligate carriers, but since they are most likely heterozygous, they will not express the trait. These female carriers however can transmit the trait to their offspring, resulting in 50% of the daughters being a carrier, and 50% of the sons being affected. Therefore, affected males can transmit an X-linked recessive disease to 50% of their grandsons through their obligate carrier daughters. In X-linked dominant diseases, an excess of affected females exists in pedigrees of the disease, and again male-to-male transmission is impossible. However, all daughters of an affected male will have the disease, since the dominant trait will be expressed in the heterozygous females. Since both sexes in the offspring of these females have an equal chance of obtaining the X-chromosome carrying the mutant allele, half off the offspring of both sexes will be affected.

Although these rules are strait and in concordance with the inheritance of sex-linked traits in *Drosophila,* several important exceptions apply to humans. Dosage compensation in *Drosophila* is achieved by up-regulation of gene expression from the single X-chromosome in males^30^. In human, however, one of the two X-chromosomes in females is silenced, and since XCI is basically random in every cell, females are a mosaic of cells in which either the maternal or paternal X-chromosome is inactivated^2^. In *Drosophila,* a recessive trait on the X-chromosome is really behaving recessively, since both X-chromosomes in a cell are active and thus being expressed. However, in women, this is not the case. If both X-chromosomes have an equal chance of being inactivated, in half of the cells the recessive allele will be inactivated and hence not expressed, whereas in the other half the recessive allele will be present on the active X-chromosome (Xa). Thus even a recessive trait will be uniquely expressed in half of the cells of women, therefore making it dominant at the cellular level, and should thus result in a phenotype similar to that in hemizygous males. Indeed, far more female carriers of an X-linked disease present with clinical symptoms^31,32^, which cannot be explained by simplified rules of dominant and recessive X-linked inheritance as obtained in the *Drosophila* model, deduced at a time that knowledge on differences between human and *Drosophila* dosage compensation was lacking^33^. In agreement with this, a systematic review of 32 X-linked diseases has shown that, as expected based on the X-linked pattern of inheritance, the severity and penetrance, which is the frequency with which a genotype manifest itself in a given phenotype, are higher in males than in females carrying an X-linked disease mutation, whereas the severity and penetrance are highly variable in heterozygous females, varying from high, to intermediate and low between different disorders, but also in females being affected with the same disease^33^. Thus apparently, for X-linked diseases, classifications as recessive and dominant are misleading, and can better be described as X-linked^33^.

## X-CHROMOSOME INACTIVATION MOSAICISM AND SKEWING AS MODIFIERS OF X-LINKED HUMAN DISEASE IN FEMALES

In complex organisms like humans, different cell types have to work together to obtain normal physiology. As a XCI consequence, and the clonal propagation of the inactivated Xi state, women are a mosaic of two cell populations, with either the maternally or the paternally inherited X-chromosome being inactivated^3^. Both populations of cells make up all organs, and despite the mixing of both different cell populations, a proper interaction is required for development and physiology. Different forms of communication are present between cells, whether it is through direct cell-cell contacts or secreted factors, which most of the time result in metabolic cooperation between cells^32^. In light of X-linked disease, these interactions can have crucial consequences for the outcome of the disease severity for female carriers of heterozygous mutations^34^. If an X-linked mutation results in the absence of a certain protein, a cell which has inactivated the wild-type, X-chromosome will experience problems from the protein absence, possibly resulting in growth disadvantage of that cell, or even cell death, which will result in selection against mutant cells. However, if the protein or signal can be acquired from cells in the surrounding of the mutant cells, for example by uptake through endocytosis or gap-junctions, the deficiency of mutant cells can be hidden, and function might be restored (**Figure 2A**). This form of metabolic cooperation between cells seems to be true for a variety of X-linked metabolic diseases, including Lesch-Nyhan syndrome, Fabry disease and Hunter syndrome which manifest themselves with severe phenotypes in affected males, but only present mild symptoms in heterozygous females. In iduronate sulfatase deficiency (Hunter syndrome, OMIM: 309900), large amounts of mucopolysaccharides accumulate in mutant cells, resulting in bone abnormalities, deafness and enlarged organs in affected males. In female heterozygote carriers of identical mutations, iduronate sulfatase is transferred from wild-type to mutant cells, and hence accumulation of harmful products does not occur in most of these women, resulting in minimal presentation or absence of disease symptoms^35^. In Fabry disease (OMIM: 301500), another lysosomal storage disease which results in accumulation of glycosphingolipids in blood and lysosomes of most cells, causing clotting of blood vessels and tissue damage, females also have milder symptoms, since the *α*-galactosidase A enzyme can transfer from unaffected to mutant cells, although at lower efficiency compared to iduronate sulfatase. In Lesch-Nyhan syndrome (OMIM: 300322), males have hypoxanthine phosphoribosyltransferase (HPRT) deficiency, causing a defect in recycling of purine bases, resulting in mental retardation, cerebral palsy, self-destructive biting behavior and joints accumulation of uric acid. Heterozygous female carriers are often unaffected, because inosinate, a product of the HPRT metabolic reaction, is transferred through gap-junctions. This transport is possible in fibroblasts, resulting in fibroblast populations consisting of both wild-type and surviving mutant cells, but in circulating blood cells, where gap-junctions are absent, this form of rescue cannot occur, resulting in selection against cells which express the mutant allele^31^. Most likely, these cells are gradually outcompeted by wild-type cells, which will have a proliferative advantage over mutant cells^34^.

**Figure 2.**
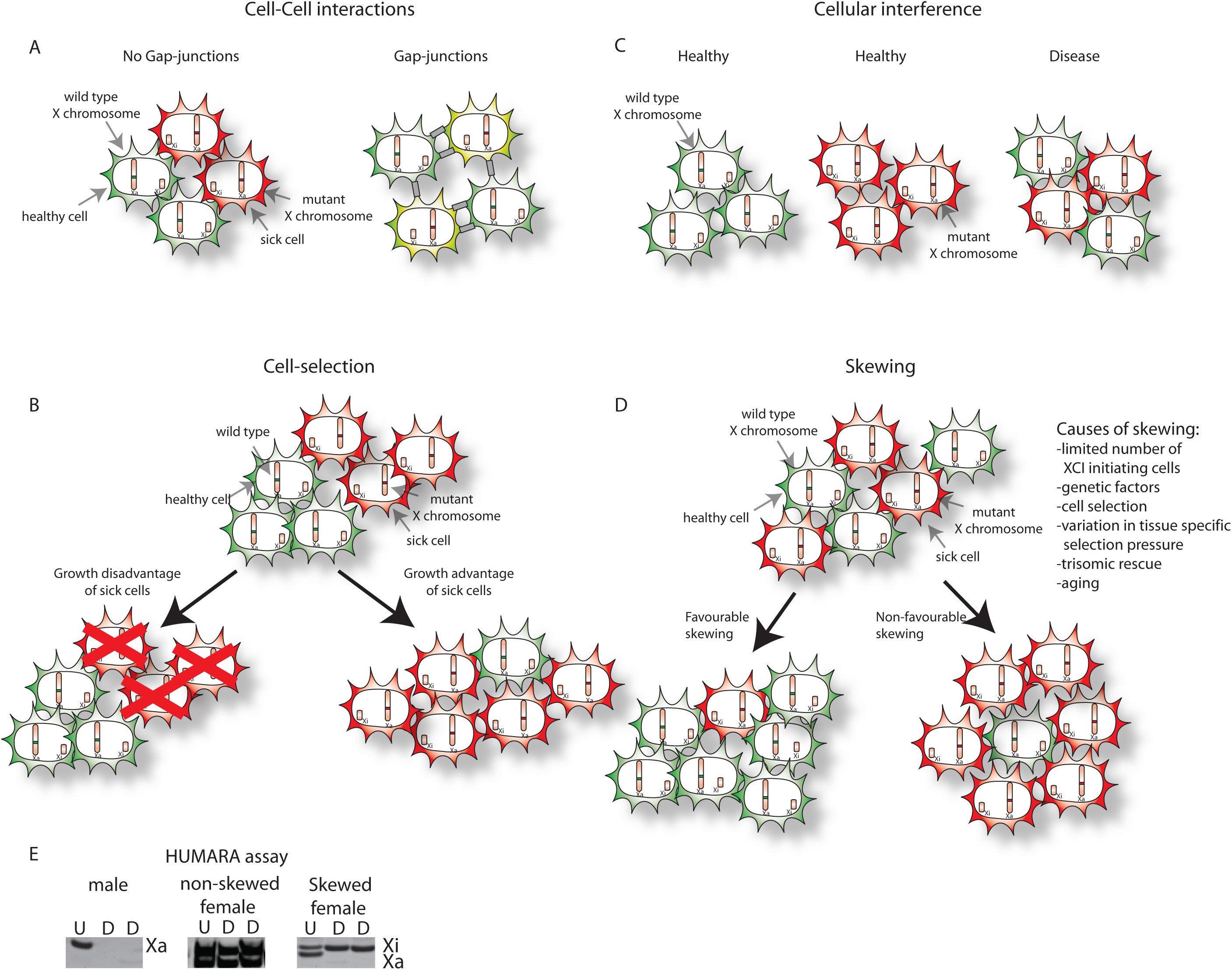
How X-inactivation can influence the disease phenotype in females. **A)** When mutations on the active X-chromosome are present, cells which have inactivated the wild-type X-chromosome will express the mutant copy of a gene, and will hence experience an absence of a functional protein. This will usually result in a cell malfunction, but this is not the case when the protein of interest can be exchanged between cells through gap-junctions. **B)** Mutations expressed on the active X-chromosome might result in a growth advantage or disadvantage of the cells, resulting in a shift in the populations of both initial cell types. **C)** In some cases, mutations of an X-linked gene do not result in a phenotype, when all cells present express either the mutant or the wild-type copy of the gene. However, when a mixed population of cells is present, cell-cell interactions result in a phenomenon called cellular interference, resulting only in a phenotype when a heterogeneous population of cells is present. **D)** Although X-chromosome inactivation results in either the silencing of the paternal or maternal X-chromosome, the ratio between both cell types is not always 50:50. Deviation from this ratio is called X-chromosome inactivation skewing, and might result in a more favourable or non-favourable disease phenotype in females affected. Skewing is caused by several mechanisms, including initiation of X-chromosome inactivation in a limited pool of progenitor cells, genetic factors, cell selection mechanisms, which might be tissue specific, aging and more peculiar processes like trisomic rescue in early embryos. **E)** X-chromosome inactivation skewing analysis in females using the HUMARA assay. To determine the degree of X-Chromosome inactivation skewing, a frequently used method is the Human Androgen Receptor assay (HUMARA), which makes use of the presence of a polymorphic repeat in the *androgen receptor* and methylation of the inactive X-chromosome. Prior to digestion with a methylation-sensitive enzyme (undigested, U) PCR analysis will show a single allele in males (left panel), and two alleles in females (when polymorphism is present). Upon digestion with a methylation-sensitive enzyme (digestion, D), only the allele which is inactivated, and hence methylated, will be amplified. Therefore in males, upon digestion, no band is detectable. When X-chromosome inactivation is random in females, 50% of cells will methylate the one allele, whereas 50% of cells will methylate the second allele. Therefore, in a non-skewed female, the same bands will be seen upon digestion as in the undigested sample (middle panel). Upon complete skewing (right panel), only a single allele will be detected.

Most of the time, the presence of an X-linked mutation results in growth disadvantage of cells expressing the mutated allele^36^ but there are exceptions (**Figure 2B**). In adrenoleukodystrophy (OMIM: 300100), were mutations in the *ABCD1* gene result in a non-functional ALDP protein, cells expressing the mutated protein proliferate faster^37^. ALDP is involved in transport of fatty acids into peroxisomes, were they are being oxidized in the beta-oxidation pathway, as a means of energy production. When ALDP function is ablated in males, intracellular fatty acid accumulation causes loss of function of the adrenal glands, and progressive destruction of myelin in the central nervous system. Several variations of the disease exist, which affect only the adrenal glands, or show a phenotype also in the central nervous system or even the spinal cord (adrenomyeloneuropathy). Heterozygous females are often symptom free at the beginning of their life, but during aging they can develop spinal cord symptoms, which indicates that mutant cells gradually take over, due to an unexplained growth advantage^38,39^.

Interestingly, not always does cell communication results in amelioration of an X-linked disorder and in some cases it might even worsen the phenotype in heterozygous females (**Figure 2C**). This is the case for *EFNB1* gene mutations which cause craniofrontonasal syndrome (OMIM: 304110)^40^. The absence of functional ephrin B1 protein, a signaling molecule involved in many developmental processes, results in premature closure of the coronal suture (craniosynostosis), and thus aberrant skull development, only in females. Males with mutations in EFNB1 do not develop these skull deformations, because other redundant signaling molecules are able to replace the function of ephrin B1. In case of heterozygous females, where XCI has resulted in a mixture of cells which are either positive or negative for ephrin B1, such a redundant signaling pathway is disrupted by a mechanism called cellular interference^41^. A similar mechanism has been proposed in an affected boy, carrying a mosaic of a supernumerary small ring X-chromosome harboring EFNB1^42^, and in affected males harboring somatic mosaicism with an EFNB1 mutant cell line^43^.

Although XCI in principal results in two different cell populations, and this can have profound effects on the manifestation of X-linked diseases in heterozygous females, another layer of complexity is added by the fact that the ratio of the two cell lines, with either the maternal or paternal X-chromosome being inactivated, is not always 50:50. In fact, studies have shown that in large populations of healthy women, the ratio between both cell types follows a bell-shaped distribution, with a mean of 50:50 and extremes approaching 100:0 or 0:100^44^. Just by chance, a proportion of women will have a deviation from the 50:50 ratio and this deviation is called XCI-skewing (**Figure 2D, E**). How can skewing be explained? Several factors might contribute, and most likely one of the most important factors will be the presence of a limited amount of cells at the moment that XCI is initiated. Even when every cell has a 50% chance of inactivating the one or the other X-chromosome, when the starting pool of cells is small, just by chance the majority of cells might choose to inactivate for example the paternal X-chromosome. Although it is not exactly known how many cells are present at the moment that XCI is initiated in the human embryo, it is estimated that the number of cells which commence to initiate XCI is rather low, with estimates varying between 10 to 100 cells ^25,45^. Hence, with such a small pool of progenitor cells, it seems likely that the stochastic nature of the process plays an important role in determining the XCI-fate of the X-chromosomes present.

XCI-skewing likely is also genetically determined^46^. In mice, distortions from random-XCI are observed when several inbred mouse strains are intercrossed, and since these deviations are not caused by cell selection, they are referred to as primary non-random-XCI. This has resulted in the classification of mouse strains according to differences in strength of the X controlling element (Xce), a genetically defined locus which affects skewing^47^. Although the identity of this locus is still unknown, genetic mapping studies have indicated that it lies downstream of *Xist*^48^. Alternatively the amount of skewing might correlate with the presence of SNPs in XCI-regulatory elements, including the *Xist* promoter or gene itself, or other regulators, like XCI-activators and -inhibitors, or their binding places or cis-acting landscape. In agreement with this, SNPs have been found in the human *XIST* promoter, correlating with skewing^49,50^. In both cases, a CTCF binding site seems to be affected, by a nucleotide change located at −43 bp of the *XIST* transcriptional start site. Whether an XCE exists in human is unclear^51^. Several families have been identified in which extreme XCI-skewing is inherited, but this extreme skewing could also be explained by the presence of unidentified X-linked disease alleles which are being selected against^52^ when affected cells have a proliferative disadvantage, or die, as discussed above. Therefore, skewing could also be caused by X-linked mutations affecting cell viability, followed by negative selection, or sometimes even positive selection, as in the case of adrenoleukodystrophy. An even more extreme form of this secondary non-random-XCI, is found in so-called trisomic zygote rescue, and might also be causative in the development of skewing. Some cases have been described, in which placental tissues consist of a high proportion of trisomic cells, whereas the resulting fetus consist of diploid cells, which have a severely skewed XCI-phenotype^53^. It has been hypothesized that such fetuses may survive, since the initial zygote, which must have been trisomic as well, lost the supernumerary chromosomes. Since such a lucky event will only occur in a minority of cells in such an otherwise lethal embryo, even more cells might have been lost from the already small pool of cells which initiate XCI, resulting in extreme skewing observed in the girl born. Such a reduction in progenitor pool size might also underlay the observed more extreme skewing in monozygotic twins compared to singlets^54,55^, although not all studies agree on this^45,56^.

The selection pressure for or against cells carrying an X-linked mutation does not seem to be identical in all tissues or at all developmental stages. For example, it has been found that several X-linked diseases only result in tissue-specific skewing. In X-linked agammaglobulinemia (Bruton’s agammaglobulinemia, OMIM: 300300), caused by mutations in BTK, mutant B-cells are unable to mature, and are thus absent in the peripheral blood system of heterozygous females, where wild-type B-cells are selected for^57^. The mutant B-cell progenitors however can be found in bone marrow, indicating that different stringencies of selective pressure are maintained in different tissues. Also other cell types, where *BTK* is not expressed, will most likely be confronted with an absence of selection pressure against mutant cells. The same tissue-specific skewing can be found in other hematological X-linked diseases (**Supplementary Table 1**), including X-linked severe combined immunodeficiency (X-SCID, OMIM: 300400) and Wiskott-Aldrich syndrome (OMIM: 301000)^58^, and extreme skewing has even been used to identify carriers of disease mutations prior to genetic testing of the disease genes^59^.

**Table 1.** X-linked disorders associated with favorable skewing and minimal disease in females.

| **Disease** | **Gene** | **Gene location** | **OMIM** | **Phenotype in males** | **PMID skewing in females** |
| --- | --- | --- | --- | --- | --- |
| **Agammaglobulinemia** | *BTK* | Xq21.33-q22 | 300300 | Decreased number of peripheral B-cells, decreased antibody levels and recurrent infections | 7506482, 9633977 |
| **Alpha-thalassemia / mental retardation syndrome** | *ATRX* | Xq21.1 | 301040 | Dysmorphic features, mental retardation, hemoglobin H disease, epilepsy | 1415255, 16100724 |
| **Barth syndrome** | *TAZ* | Xq28 | 302060 | Cardiomyopathy, neutropenia, skeletal myopathy and growth delay | 9792874 |
| **Dyskeratosis congenita** | *DKC1* | Xq28 | 305000 | Premature aging, abnormal skin pigmentation, nail dystrophy, leukoplakia, progressive bone marrow failure | 9311754, 9042917, 9310472 |
| **Dystonia-deafness-optic neuropathy syndrome** | *DDP* | Xq22.1 | 304700 | Progressive form of deafness, visual disability leading to cortical blindness, dystonia, fractures, mental deficiency | 8826445 |
| **Hypohidrotic ectodermal dysplasia with immunodeficiency** | *NEMO* | Xq28 | 300291 | Hypomorphic mutations resulting in skin lesions and recurrent infections | 16333836 |
| **MECP2 duplication syndrome** | *MECP2* | Xq28 | 300260 | Intellectual deficits, autism, muscular hypotonia, frequent recurrent infections and mild dismorphic features | 16080119 |
| **Severe combined immunodeficiency syndrome** | *IL2RG* | Xq13 | 300400 | Increased susceptibility to infections, low antibody levels, thymus atrophy | 1395081, 8167715, 1550118, 9633977 |
| **Wiskott-Aldrich syndrome** | *WASP* | Xp11.23-p11.22 | 301000 | Thrombocytopenia with small-sized platelets, eczema, recurrent infections | 6101415, 7437512, 3263154, 2043768 |

Skewing is not only influenced by tissue specificity, but also by aging. Interestingly, many studies have found that skewing is more prevalent in elderly women^60^. Certain X-linked diseases only develop in elderly women due to age-related skewing. Examples include sideroblastic anemia (OMIM: 300751)^61^ and X-linked hemolytic anemia (OMIM: 305900) caused by G6PD deficiency^62^. The reason for this is unclear, but might involve stochastic clonal loss^63^, or genetic selection, potentially due to subtle, hidden mutations or SNPs affecting gene function or expression of X-linked genes, which manifest themselves gradually in growth advantage or disadvantage^64^.

What is clear from the issues discussed above is that the variability of symptoms in heterozygous female carriers of an X-linked disease can be influenced by many factors. Genetic and stochastic factors will first determine the mosaic distribution of cells in embryonic tissues having inactivated the maternal or paternal X-chromosome. This immediately results in a mosaic distribution of mutant and wild-type cells, through the growing tissues. The effect of the mutation itself on the cell viability and growth of the mutant cells will further result in a complicated interplay between wild-type and mutant cells, either favoring wild-type cells when mutations result in a negative outcome for the affected cells, or favoring the mutant cells when mutations are advantageous. When mutations affect cell-autonomous proteins and pathways, a different outcome can be expected compared to the situation in which mutant cells can be rescued by wild-type proteins obtained from neighboring cells. Thus not only does skewing influence the outcome of genetic disease, also the opposite is true; namely, due to X-linked disease the ratios between cells having inactivated the wild-type or mutant X-chromosome will differ, dependent on the effect of the mutation. However, since females have two X-chromosomes, favorable XCI-skewing, either primary or secondary due to selection, can thus often result in a better outcome of X-linked diseases compared to males, which suffer from their higher vulnerability for X-linked diseases due to the presence of the hemizygous X-chromosome, lacking a backup allele for X-linked loci. **Table 1** summarizes several diseases in which favorable skewing has been found in heterozygous carriers of X-linked disease, presenting themselves with minimum symptoms compared to males.

## WHEN MOSAICISM CAUSES A DISADVANTAGE: EMERGENCE OF MANIFESTING HETEROZYGOTES

Not always does the mosaic expression or completely skewed XCI result in a benefit for a woman. This can occur when extreme skewing results in inactivation of one X-chromosome, maybe because of selection against a mutant allele, but unfortunately also the Xa harbors a genetic defect. This disease allele, which would otherwise maybe be selected against, or the disease outcome would benefit from a mosaicism, will now be active in all cells, causing disease which is otherwise not observed in heterozygous female carriers. Such a case has been described in which a female inherited a maternal mutation in the incontinentia pigmenti gene *NEMO* (OMIM: 308300), and a paternal mutation in the *F8* gene causing hemophilia A (OMIM: 3067000)^65^. The *NEMO* mutation is detrimental for cells, and thus strongly selected against, rendering all cells with the wild-type incontinentia pigmenti gene active. Unfortunately for this patient, the X-chromosome containing the wild-type *NEMO* gene also contains the mutant *F8* gene, therefore resulting in a bleeding disorder.

Many cases of these so-called manifesting heterozygotes have been described (**Supplementary Table 2**), and some of these have even been crucial for the identification of the disease alleles involved. The reason for extreme skewing which results in the manifestation of the X-linked disease can be variable, ranging from chance to genetic causes, as discussed above. In the case of a girl presenting with Wiskott-Aldrich syndrome (OMIM: 301000)^66^, the affected girl, her mother and grandmother showed extreme skewing, pointing to a genetic cause. The mutant Wiskott-Aldrich syndrome was located on the Xa, causing the disease, which is normally not seen in females due to XCI of the X carrying the disease allele. Other women with Wiskott-Aldrich syndrome have been described, where XCI-skewing resulted in 100% of cells with the mutated allele on Xa^67^, or absence of skewing against the mutant allele was observed^68^. Other similar examples includes affected females with hemophilia B (OMIM: 306900), myotubular myopathy (OMIM: 310400), X-linked hemolytic anemia (OMIM: 305900), X-linked thrombocytopenia (OMIM: 313900), ATRX syndrome (OMIM: 301040) and Fabry disease (OMIM: 301500), where also extreme XCI-skewing towards inactivation of the wild-type X-chromosome was found (**Supplementary Table 2**).

**Table 2.**
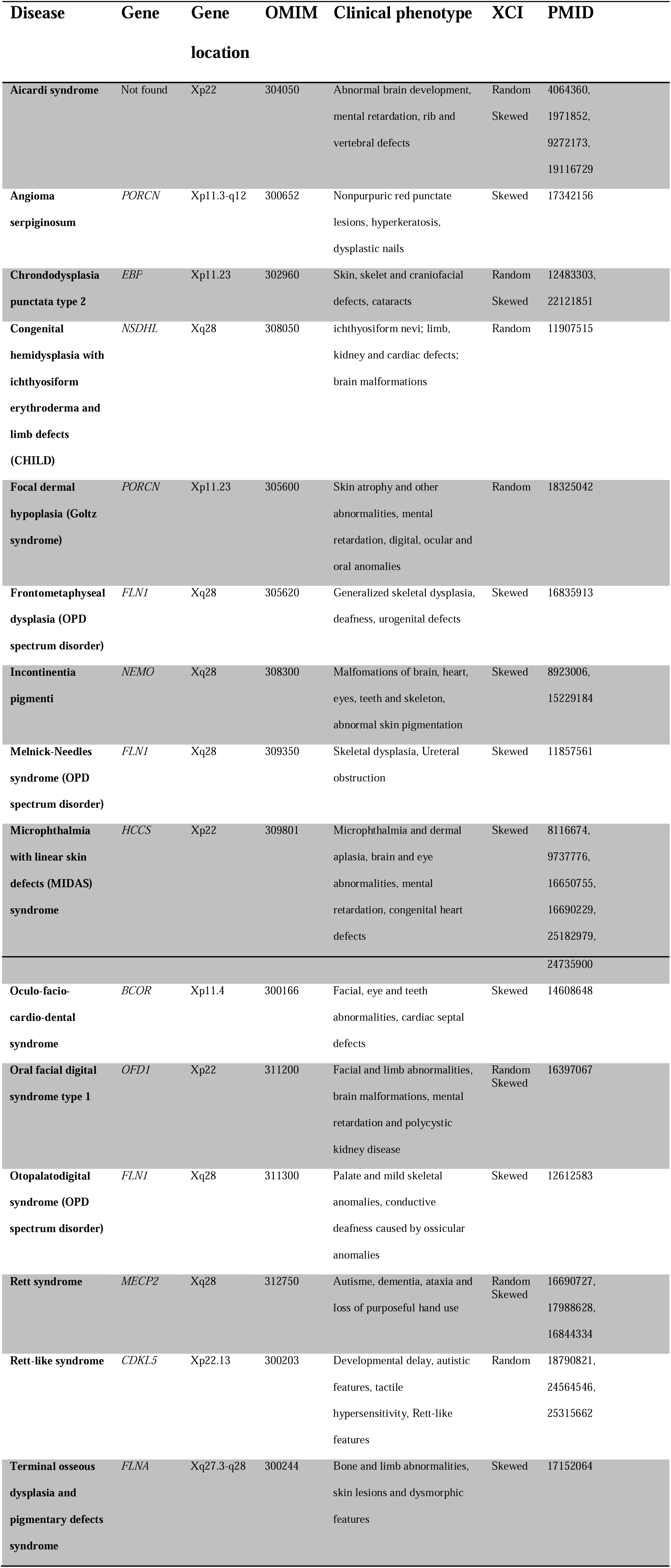
X-linked Disorders with reduced viability in males.

| Disease | Gene | Gene location | OMIM | Clinical phenotype | XCI | PMID |
| --- | --- | --- | --- | --- | --- | --- |
| **Aicardi syndrome** | Not found | Xp22 | 304050 | Abnormal brain development, mental retardation, rib and vertebral defects | Random<br>Skewed | 4064360,<br>1971852,<br>9272173,<br>19116729 |
| **Angioma serpiginosum** | *PORCN* | Xp11.3-q12 | 300652 | Nonpurpuric red punctate lesions, hyperkeratosis, dysplastic nails | Skewed | 17342156 |
| **Chondrodysplasia punctata type 2** | *EBP* | Xp11.23 | 302960 | Skin, skelet and craniofacial defects, cataracts | Random<br>Skewed | 12483303,<br>22121851 |
| **Congenital hemidysplasia with ichthyosiform erythroderma and limb defects (CHILD)** | *NSDHL* | Xq28 | 308050 | ichthyosiform nevi; limb, kidney and cardiac defects; brain malformations | Random | 11907515 |
| **Focal dermal hypoplasia (Goltz syndrome)** | *PORCN* | Xp11.23 | 305600 | Skin atrophy and other abnormalities, mental retardation, digital, ocular and oral anomalies | Random | 18325042 |
| **Frontometaphyseal dysplasia (OPD spectrum disorder)** | *FLNI* | Xq28 | 305620 | Generalized skeletal dysplasia, deafness, urogenital defects | Skewed | 16835913 |
| **Incontinentia pigmenti** | *NEMO* | Xq28 | 308300 | Malfomations of brain, heart, eyes, teeth and skeleton, abnormal skin pigmentation | Skewed | 8923006,<br>15229184 |
| **Melnick-Needles syndrome (OPD spectrum disorder)** | *FLNI* | Xq28 | 309350 | Skeletal dysplasia, Ureteral obstruction | Skewed | 11857561 |
| **Microphthalmia with linear skin defects (MIDAS) syndrome** | *HCCS* | Xp22 | 309801 | Microphthalmia and dermal aplasia, brain and eye abnormalities, mental retardation, congenital heart defects | Skewed | 8116674,<br>9737776,<br>16650755,<br>16690229,<br>25182979, |
|  |  |  |  |  |  | 24735900 |
| **Oculo-facio-cardio-dental syndrome** | *BCOR* | Xp11.4 | 300166 | Facial, eye and teeth abnormalities, cardiac septal defects | Skewed | 14608648 |
| **Oral facial digital syndrome type 1** | *OFD1* | Xp22 | 311200 | Facial and limb abnormalities, brain malformations, mental retardation and polycystic kidney disease | Random Skewed | 16397067 |
| **Otopalatodigital syndrome (OPD spectrum disorder)** | *FLN1* | Xq28 | 311300 | Palate and mild skeletal anomalies, conductive deafness caused by ossicular anomalies | Skewed | 12612583 |
| **Rett syndrome** | *MECP2* | Xq28 | 312750 | Autisme, dementia, ataxia and loss of purposeful hand use | Random Skewed | 16690727,<br>17988628,<br>16844334 |
| **Rett-like syndrome** | *CDKL5* | Xp22.13 | 300203 | Developmental delay, autistic features, tactile hypersensitivity, Rett-like features | Random | 18790821,<br>24564546,<br>25315662 |
| **Terminal osseous dysplasia and pigmentary defects syndrome** | *FLNA* | Xq27.3-q28 | 300244 | Bone and limb abnormalities, skin lesions and dysmorphic features | Skewed | 17152064 |

Another frequent cause of manifesting heterozygotes are X-chromosome translocations^69^. In translocations, two breaks occur, one in the X-chromosome, and another one on an autosome, after which the broken parts are joined together. Generally, either a balanced or unbalanced type of X-to-autosome translocations can be distinguished. In unbalanced X-translocations, an additional part of the X-chromosome is present, linked to a piece of autosome, representing a partial trisomy of that autosome. In general, trisomies of autosomes are not tolerated, except for partly viable trisomies of chromosome 13, 18, 21 (Patau, Edwards, and Down syndromes, respectively) which nevertheless result in severe health problems. An unbalanced X-autosome translocation will only be viable if the translocated piece of X-chromosome is able to undergo inactivation and to silence the attached piece of autosome by spreading of XCI. Therefore the translocated piece of X-chromosome must at least contain the XIC, located at Xq13. If this is not the case, this will most likely result in a non-viable situation, since cells will be confronted with a higher dosage of autosomal genes. If, in the case of an unbalanced translocation, there is no duplicated, extra part of autosome present, silencing of the translocated X-chromosome might result also in silencing of the attached autosomal segment, resulting in monosomy of autosomal genes, which is also likely to be lethal. In case of balanced translocations, the situation is different^70^. Here, the total amount of chromosome parts is the same as in a normal genome, but one piece of X-chromosome is translocated to a part of an autosome, and the other two broken parts are also joined together. Since these cells carry two X-chromosomes, XCI will be initiated. However, if inactivation is initiated on the X-autosome translocation product, this leads to silencing of autosomal genes, leading to a monosomy of the autosomal part of the fusion chromosome. Since most autosomal monosomies are incompatible with life, cells having inactivated the translocated chromosome will be selected against. Therefore, in a balanced translocation, the wild-type X-chromosome is always inactivated, leaving the translocation products active, to result in the best genetic balance. Hence, most of the carriers of balanced translocations have a normal phenotype, although they might be sterile^70^. However, in the process of translocation break and repair, parts of the X-chromosome might get lost, or the breakage can disrupt gene coding or regulatory sequences. Since the wild-type X-chromosome in these balanced X-to-autosome translocations is always inactivated, the disrupted gene will be on the active (parts of the) X-chromosome(s), and therefore can result in a manifesting disease in females. Examples of this include girls with hypohidrotic ectodermal dysplasia (OMIM: 305100), Lowe oculocerebrorenal syndrome (OMIM: 309000), Simpson-Golabi-Behmel syndrome (OMIM: 312870), Hunter syndrome (OMIM: 309900) and hyper IgM immunodeficiency syndrome (OMIM: 308230). Such X-to-autosome translocations have also been helpful in the mapping of several X-linked disease genes, including genes underlying Duchenne muscular dystrophy (OMIM: 310200), oligophrenin-1 syndrome (OMIM: 300486), Aarskog-Scott syndrome (OMIM: 305400), incontinentia pigmenti (OMIM: 308300) and chronic granulomatous disease (OMIM: 306400) (**Supplementary Table 2**).

## DISEASES WHERE FEMALES LIKELY BENEFIT FROM THEIR MOSAICISM: MALE LETHAL DISEASES AND DISEASES WITH REDUCED VIABILITY OF MALES

A severely detrimental mutation on the single X-chromosome will result in male lethality, so that no males are born carrying such a disease allele. Indeed, several X-linked disorders have been described which result in male lethality, or highly reduced male viability, and therefore these diseases are only found in females (**Table 2**). This group of disorders is characterized by symptoms in heterozygous females, and they were originally classified as being X-linked dominant. However, symptoms amongst affected females are variable, with some patients presenting the full disease phenotypes, whereas others present as asymptomatic carriers. In most instances, XCI-skewing and mosaicism are responsible for the different phenotypes observed, and explain that many affected females are alive despite carrying a disease allele which is lethal in males. Indeed, in the majority of these diseases, favorable skewing is associated with reduced symptoms. In case of Rett syndrome (OMIM: 312750), where mutations in the MECP2 gene cause autism, ataxia, tremors, loss of purposeful hand use and dementia, XCI is often random amongst affected females^71^. However, some sporadic, unaffected female carriers of the mutation, show preferential inactivation of the disease allele^72^, and favorable skewing has been observed amongst milder affected females^73^. For Rett syndrome selection against cells that have inactivated the wild-type X chromosome does not happen, and the phenotype is therefore directly related to skewing effects originating as a consequence of the XCI process (primary non-random-XCI), or secondary effects due to selection in favor of other traits (secondary non-random-XCI). In another male lethal neurodevelopmental disorder, called microphthalmia with linear skin defects syndrome (OMIM: 309801) caused by mutations in the *HCCS* gene, skewed XCI has been observed in the majority of cases^74^. Here it seems that the presence of severe skewing is the only reason that female patients are alive, as skewing has been reported in all heterozygous females Apparently, in HCCS mutant cells secondary cell selection in the majority of tissues has occurred against cells with the mutant allele on the Xa. The efficiency with which this process occurs in different tissues might explain the variable phenotypes observed between different affected females harboring identical mutations. Besides favorable skewing, another advantage for females might be the possibility of genes escaping XCI. For both incontinentia pigmenti (OMIM: 308300) and oral-facial-digital syndrome type I (OMIM: 311200) it has been found that these genes escape XCI in humans^75^. For the latter syndrome, this might explain why human females carrying an *OFD1* mutation are alive, whereas mice with a heterozygous *Ofd1* knockout allele die, because *Ofd1* is not escaping XCI in mouse^76^. Escape most often is incomplete and might be tissue-specific, and this might explain why XCI-skewing is still observed in some cases of these syndromes^77,78^. A similar advantage due to escaping XCI has been proposed for the X-linked histone H3K27me3 demethylase UTX in human T-cell acute lymphoblastic leukemia, where detrimental mutations leading to leukemia were only found in male patients but not in females^79^.

## X-LINKED DISEASE IN FEMALES NOT RELATED TO X-CHROMOSOME INACTIVATION OR SKEWING

As is clear from the discussion above, XCI can have a great influence on the phenotype and severity of X-linked diseases. However, not all female diseases due to X-linked mutations are necessarily influenced by XCI. Besides what is seen in manifesting heterozygotes due to unfavorable skewing, females might also experience symptoms of an X-linked disease just due to the fact that they inherited two mutant alleles for the same disease. Examples of homozygotes for X-linked diseases include females with hemophilia^80^, congenital X-linked nystagmus (OMIM: 310700)^81^, X-linked recessive ichthyosis (OMIM: 308100)^82^, hypophosphatemic rickets (OMIM: 307800)^83^ and Becker muscular dystrophy (OMIM: 300376)^84^, amongst others. The chance of inheriting two X-linked recessive alleles is low, except for genetically homogenous families or families where X-linked diseases are known to occur on both parental sides. In addition, homozygous mutations for X-linked genes can also be caused by *de novo* mutations^85^, or in rare causes of uniparental disomy, where both X-chromosomes present are derived from the same parent^86^. Also, manifestations of X-linked diseases in females sometimes discover a hidden, or not yet diagnosed form of Turner syndrome, in which a female has only one X-chromosome, thus rendering a Turner female prone to the same risk of developing an X-linked disease as a hemizygous male^87,88^. Even more rarely, the presence of an X-linked disease has discovered cases of dysregulated sex-determination, where affected females turned out to have a male karyotype^89,90^.

The benefits or disadvantages of mosaicism related to X-linked gene expression may not always be related to XCI. For example, rare cases of mosaicism may be caused by fusion of two distinct zygotes, or due to placental transport of cells between twins^91^. Such chimaeras might display mosaicism, in which one population of cells harbors an X-linked mutation, whereas the other carries a wild-type allele. Also mutations occurring during postzygotic stages of development will result in the fact that not all cells of the body will harbor a mutation. The later in embryonic development the mutation occurs, the fewer cells will be affected^92^. In such cases, it is also possible that the mutation does not affect the germ line, which is important for counseling issues and the risk of transmitting the disorder to the offspring

## CONCLUSION AND OUTLOOK

It has become clear in recent years that XCI is not only a fascinating biological process occurring during early female embryonic development, but also an important modifier of X-linked disease in females. The accompanying online-only discussion further elaborates on this, by providing an extensive review of hematological, immunological, neurological, muscle, metabolic, renal, dermatological, and ocular diseases affected by XCI, and the influence of this process on cancer, pregnancy loss and structural and numerical disorders of the X-chromosome. Whereas the ancient opinion that X-linked disease is only relevant for males has become a subject of change, which is emphasized by the large number of reports linking XCI-skewing to the phenotypes in females discussed in this review, in the next step it will be crucial to link the fundamental knowledge obtained by basic research on the regulation of XCI-initiation to future clinical implications. For example, it is highly likely that currently non-understood effects of XCI-initiation in females explain why relatively fewer females are born compared to males. In all Western civilizations, at present the birth ratio is around male: female = 1.05:1.0, as exemplified for the Netherlands (**Supplementary Figure 1**). Whereas at the moment of fertilization, an equal ratio between both sexes is found^93^, and during recognized pregnancy and during postnatal development more males are dying (most likely reflecting the male disadvantage due to the hemizygous X-chromosome)^94^, apparently a large number of female embryos is lost around the time of implantation in the uterus^95^. It is that around that time, the only significant difference known between both sexes is the XCI-process, and hence it is likely that effects of this process, as they might for example occur due to stochastic XCI-initiation on too many or less X-chromosomes as observed in differentiating mouse ES cells and embryos, are contributing to this observation. Our fundamental insights in XCI might help to develop targeted X-reactivation methods, which could be used to treat heterozygous females suffering from X-linked disease due to preferential inactivation of the wild-type X-chromosome (**Figure 3**). Some first successes have been obtained by applying such ideology to the treatment of Rett syndrome^96,97^, and recent efforts to reactivate the X chromosome are promising^98^, although this strategy should also be taken with caution as this could have adverse effects as well^99^. Nevertheless, we foresee that important new breakthroughs, such as targeted X-reactivation, will follow in the near future, bridging the gap between fundamental XCI-research and the clinic.

**Figure 3.**
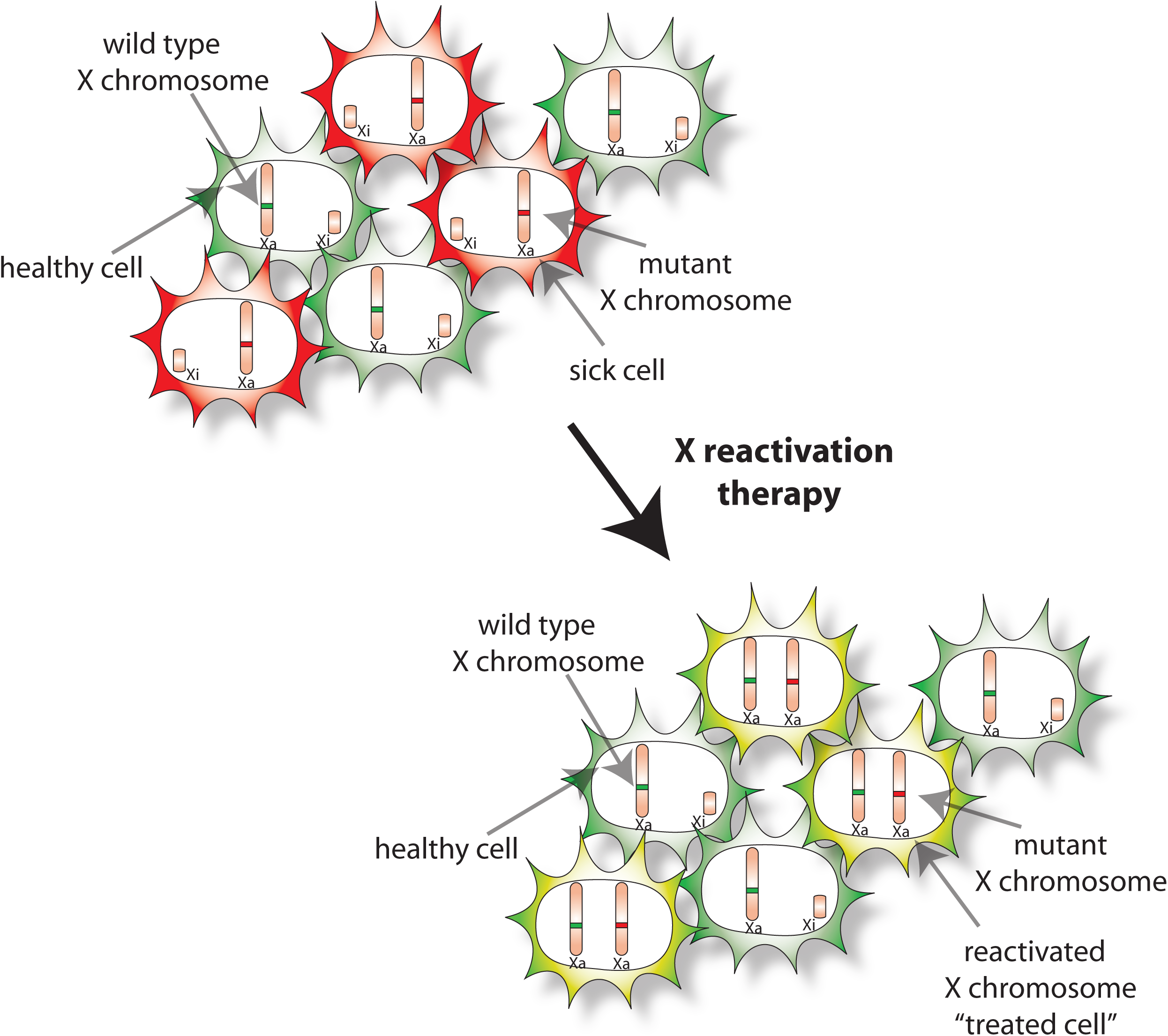
X chromosome reactivation as a potential novel treatment for X-linked disorders in females. When a wild-type X chromosome which is silenced could get reactivated, a cell will re-express a wild-type copy of the missing or non-functioning protein, which might result in an improvement of the disease phenotype.

## ACKNOWLEDGEMENTS

We would like to thank all the patients and our fellow researchers in the XCI community without whose contribution this review would not have been possible. We apologize to those whose work could not be cited due to strict space limitations. We would like to thank Robert Jan Galjaard for critically reading and valuable comments on an earlier version of this manuscript. J.G is supported by funding from the Dutch Research Council and the European Research Council. T.S.B was supported by a Niels Stensen Fellowship.

---

**Supplementary Figure 1**: Birth and child death for males and females in the Netherlands from 2004 to 2008.

*Source: Centraal Bureau voor de Statistiek (http://www.CBS.n1). retrieved 19 april 2012*

A) Table showing the total number of birth, the number of females and males and the relative ratio between both sexes born in the Netherlands during the period 2004-2008.

B) Number of still born children, without any signs of life, after a pregnancy of 22 weeks or longer, per 1000 live birth.

C) Perinatal mortality rate, defined as the number of still born children, without any sign of life, or children who died in their first 7 days of life, after a pregnancy of 22 weeks or longer, per 1000 live birth.

D) Neonatal mortality rate, defined as the number of children who died between day 1 and day 28 after birth, after a pregnancy of 22 weeks or longer, per 1000 live birth.

E) Infant mortality rate, defined as the number of children who died prior to their first 1 years birthday, after a pregnancy of 22 weeks or longer, per 1000 live birth.

**Supplementary Table 1**: X-linked primary immunodeficiencies

**Supplementary Table 2:** Examples of manifesting heterozygotes

**Supplementary Table 3**: X-linked disorders affecting the kidney

**Supplementary Table 4**: X-linked disorders affecting the skin

**Supplementary Discussion**

